# The beta-tubulin isotype TUBB6 controls microtubule and actin dynamics in osteoclasts

**DOI:** 10.1101/2021.09.17.460637

**Authors:** Justine Maurin, Anne Morel, David Guérit, Julien Cau, Serge Urbach, Anne Blangy, Guillaume Bompard

**Author notes:** These authors share senior authorship (Anne Blangy, Guillaume Bompard).

## Abstract

Osteoclasts are bone resorbing cells that participate in the maintenance of bone health. Pathological increase in osteoclast activity causes bone loss eventually resulting in osteoporosis. Actin cytoskeleton of osteoclasts organizes into a belt of podosomes, which sustains the bone resorption apparatus and is maintained by microtubules. Better understanding of the molecular mechanisms regulating osteoclast cytoskeleton is key to understand the mechanisms of bone resorption, in particular to propose new strategies against osteoporosis. We reported recently that β-tubulin isotype TUBB6 is key for cytoskeleton organization in osteoclasts and for bone resorption. Here, using an osteoclast model CRISPR/Cas9 KO for Tubb6, we show that TUBB6 controls both microtubule and actin dynamics in osteoclasts. Osteoclasts KO for Tubb6 have reduced microtubule growth speed with longer growth life time, higher levels of acetylation and smaller EB1-caps. On the other hand, lack of TUBB6 increases podosome life time while the belt of podosomes is destabilized. Finally, we performed proteomic analyses of osteoclast microtubule-associated protein enriched fractions. This highlighted ARHGAP10 as a new microtubule-associated protein, which binding to microtubules appears to be negatively regulated by TUBB6. ARHGAP10 is a negative regulator of CDC42 activity, which participates in actin organization in osteoclasts. Our results suggest that TUBB6 plays a key role in the control of microtubule and actin cytoskeleton dynamics in osteoclasts. Moreover, by controlling ARHGAP10 association with microtubules, TUBB6 may participate in the local control CDC42 activity to ensure efficient bone resorption.

## INTRODUCTION

Osteoclasts are giant multinucleated cells involved in bone resorption. Their counterparts, osteoblasts, participate in new bone formation. Both cell types are thus crucial for bone homeostasis which is highly dynamic despite bone stiffness. Furthermore, whereas an equilibrium between the activities of osteoclasts and osteoblasts is crucial for bone renewal, a pathological excess in osteoclast activity results in bone loss and may lead to osteoporosis; this is the case in number of physio-pathological situations such as from age-related sexual hormone decay, chronic inflammation and cancer (Khosla and Hofbauer, 2017). Bone resorption by osteoclasts relies on their ability to organize a specific adhesive structure called the sealing zone or podosome belt, composed of actin rich αvβ3 integrin-containing adhesive structures called podosomes (Blangy et al., 2020). The sealing zone surrounds a highly convoluted plasma membrane domain: the ruffled border, which is specifically involved in bone resorption being the siege of acid and protease secretions respectively dissolving mineral matrix and digesting bone proteins. The dynamic of this podosomal structure allows osteoclast sliding and generates series of resorption pits at the bone surface (Søe and Delaissé, 2017).

Various approaches to regulate osteoclast activity have been developed mostly focused on pathways regulating differentiation but targeting the sealing zone formation also appears an interesting alternative for the treatment of osteoporosis (Blangy et al., 2020; Mounier et al., 2020). Combining transcriptomic and proteomic approaches, we recently highlighted the β-tubulin isotype TUBB6 as important for the establishment and/or the maintenance of the podosome belt and for osteoclast activity (Guérit et al., 2020). Microtubules are dynamic polymers of α- and β- tubulin heterodimers, which oscillate between growing and shrinking phases a process known as dynamic instability. The specific regulation of MT dynamics and organization allows their adaptation to diverse cellular functions, such as cell division, motility, shape and intracellular transport. Tubulin isotypes and post-translational modifications generate a so called “tubulin code” translated into a given function through the recruitment of specific microtubule-associated proteins (Janke and Magiera, 2020). Indeed, the nature of β-tubulin isotypes was shown to affect microtubule dynamics *in vitro* (Vemu et al., 2017; Ti et al., 2018). TUBB6 was found to exert a microtubule-destabilizing effect in cycling cells (Bhattacharya et al., 2011; Bhattacharya and Cabral, 2009). Moreover, TUBB6 is expressed at low levels in most tissues (Leandro-García et al., 2010) and high expression of TUBB6 is characteristic of osteoclasts (Guérit et al., 2020). Microtubules and the actin cytoskeleton are physically and functionally intimately connected (Dogterom and Koenderink, 2019). In particular in osteoclasts, both actin cytoskeleton and microtubules are key for the formation of the podosome belt and for bone resorption (Okumura et al., 2006), but their regulatory crosstalk remains poorly understood in osteoclast (Blangy et al., 2020).

We reported recently that TUBB6 plays a key role in actin organization in osteoclasts and their resorption activity (Guérit et al., 2020). In the present study, we investigated further how TUBB6 regulates actin cytoskeleton in osteoclast, through the development of a cellular model knocked out for Tubb6 gene. We found that actin organization and bone resorption defects of Tubb6 KO osteoclasts are corelated not only with changes in microtubule but also actin dynamics. Furthermore, we developed a proteomic approach and identified proteins whose association with microtubules is affected by TUBB6.

## MATERIALS AND METHODS

### Mice, cell lines and lentivirus production

Four-week-old C57Bl/6J mice were purchased from Harlan France and maintained in the animal facilities of the CNRS in Montpellier, France. Procedures involving mice were performed in compliance with local animal welfare laws, guidelines and policies, according to the rules of the regional ethical committee. RAW264.7 cells were a gift from Kevin P McHugh (Gainesville, FL, USA). Lentiviral particles were produced at the Plateforme de Vectorologie de Montpellier (Montpellier, France).

### Osteoclast differentiation and activity

For osteoclast differentiation, RAW264.7 cells were cultured in a humidified incubator at 37 °C and 5% CO_2_, in α-MEM supplemented with 10% heat inactivated fetal calf serum (Biowest), Penicillin-Streptomycin (Life Technologies), 2 mM glutamine (Lonza) and 50 ng/mL RANKL (Peprotech). Medium was changed every other day until osteoclast differentiation at day 4 or 5. When appropriated, at day 2 of differentiation, siRNAs were transfected with siImporter (Millipore) in OptiMEM medium (Life Technologies) containing 50 ng/mL RANKL, as described previously (Touaitahuata et al., 2016), using 100 nM of the siRNAs described previously: luciferase control and Tubb6 siRNAs si1077 and si1373 (Guérit et al., 2020). Primary osteoclasts were obtained from mouse bone marrow as described previously (Vives et al., 2011; Touaitahuata et al., 2016). Briefly, non-adherent bone marrow cells from long bones of 6- to 8-week-old C57BL/6J mice were grown for 48 h in α-MEM containing 10% heat-inactivated fetal calf serum, 2mM glutamine and 30 ng/mL macrophage colony-stimulating factor (M-CSF; Miltenyi) to obtain bone marrow macrophages (BMMs); BMMs were then grown in the same medium complemented with 50 ng/mL RANKL for 5 days to obtain osteoclasts. Differentiation was performed on glass or on Apatite Collagen Complex (ACC)- coated coverslips, prepared as reported previously (Maurin et al., 2018).

To measure their activity, osteoclasts were detached from the culture plate by scrapping at day 3 of differentiation and seeded for 3 more days onto inorganic crystalline calcium phosphate (CaP)-coated multiwells (Osteo Assay Surface, Corning), eight wells per condition. For each condition, 4 wells were then stained for tartrate resistant acid phosphatase (TRAP) activity to measure osteoclast surface and 4 wells stained with von Kossa to measure CaP dissolution as described previously (Brazier et al., 2009). Wells were imaged with a Nikon SMZ1000 stereomicroscope equipped with a Nikon DXM 1200F CCD camera and the quantifications were carried out with ImageJ 1.53c software. In each experiment, osteoclast specific activity was expressed as the average area resorbed in the four wells stained with von Kossa normalized by the osteoclast surface in the four wells stained with TRAP.

### Generation of Tubb6 knock-out RAW264.7 cells

A gRNA targeting the second exon of mouse Tubb6 gene (5’-CGACCAGGCCGGAGGCTACG-3’) was designed and cloned in lentiCRISPRv2 (Addgene, 52961). Empty and Tubb6 gRNA-containing lentiCRISPRv2 were used to produce lentiviruses. RAW264.7 cells were infected with lentiviral particles and selected with 3 μg/ml puromycin 48 hours later. Resistant wild type and Tubb6 KO clones were individually picked, expanded and TUBB6 expression was monitored by immunoblot analysis using home-made polyclonal antibodies against the C-terminal end of mouse TUBB6, generated in rabbit at Eurogentec by injection of peptide GGGEINE as described (Spano and Frankfurter, 2010).

### Q-PCR

DNaseI-treated total RNA was prepared using the RNeasy Minikit (Qiagen); RNA was primed with 10-mer random primers and reverse transcription was catalyzed using Superscript II reverse transcriptase (Invitrogen) to generate cDNAs. Q-PCR was performed an Mx3000p PCR system (Stratagene) using the Platinium Taq DNA polymerase (Invitrogen) and SYBR Green I (Bio Wittaker) as described (Maurin et al., 2018). Primers used were 5′-ACAGTCCATGCCATCACTGCC-3′ and 5′-GCCTGCTTCACCACCTTCTT-3′ for Gapdh and 5′-GAACTATGTGCACCGGGACC-3’ and 5’GAGCTCGCTCAGCAGAATCC-3’ for Src and 5’-TGGAGGCCTCTCTTGGTGTC-3’ and CCACAAGATTCTGGGGACTC-3’ for Cathepsin K. The threshold cycle (Ct) of each amplification curve was calculated by Roche Diagnostics LightCycler 480 software using the second derivative maximum method. The relative amount of a given mRNA was calculated using the ΔCt method (Livak and Schmittgen, 2001).

### Immunoblot

Osteoclast lysates were prepared in lysis buffer (50 mM Tris pH7.4, 150 mM NaCl, 1% IGEPAL CA-630, 2 mM EDTA, 50 mM NaF, 10% glycerol, protease inhibitor cocktail). Protein concentrations were determined with Bradford. After resolution by SDS-PAGE, proteins were transferred to PVDF-FL membranes (Immobilon-FL Transfer Membrane) and analyzed by immunoblot. Primary antibodies were: TUBB6 (1:1000), ARHGAP10 (ProteinTech S5136AP, 1:1000), GAPDH (CellSignaling #2118, 1:5000), Sigma antibodies EB1 (E3406, 1:2500), K40-acetylated tubulin (T6793, 1:1000) and actin (A2103, 1:1000), Abcam antibodies Vinculin (ab108620, 1:5000), CLASP1 (ab108620, 1:5000) and Histone H3 (ab1791, 1:1000), total β tubulin (Developmental Studies Hybridoma Bank E7, 1:1000), TUBB5 (SAO.4G5, Santa Cruz Biotechnologies sc-58884, 1:5000). Secondary antibodies were: Goat-anti-mouse Dylight680 (Invitrogen, 35518), Goat-anti-mouse Dylight800 (Invitrogen, SA535521), Goat-anti-rabbit Dylight680 (Invitrogen, 35568), Goat-anti-rabbit Dylight800 (Invitrogen, SA535571). Signals were acquired using the Odyssey Infrared Imaging System (LI-COR Biosciences, USA).

### Immunofluorescence

All imaging was performed at Montpellier Ressources Imagerie (MRI) in Montpellier. For immunofluorescence, osteoclasts were differentiated on glass coverslips. Alternatively, osteoclasts at day 3 of differentiation were detached by scrapping and seeded for 2 more days on ACC as described previously (Maurin et al., 2018). Cells were fixed for 5 minutes in methanol at −20°C followed by 10 minute-permeabilization in 0.1% Triton X-100 in PBS in the case of endogenous EB1 labeling. Alternatively, cells were fixed for 20 minutes at room temperature in 3.2% paraformaldehyde and 10 μM Paclitaxel (Sigma) in PHEM (60 mM PIPES, 25 mM HEPES, 10 mM EGTA, 4 mM MgSO4, pH 6.9) and cells were then permeabilized with 0.1% Triton X-100 in PBS for 1 minute. After blocking for 1 hour with 2% BSA in PBS, cells were processed for immunofluorescence. Primary antibodies used were: TUBB6 (1:1000), α-tubulin (1:2000, Sigma T5168) or α-tubulin (1:200, YOL3/4 Santa Cruz Biotechnologies sc53030), ARHGAP10 (1:200), EB1(BD 610534, 1:500) in 2% BSA in PBS. Secondary antibodies used were from Invitrogen: donkey anti-mouse Alexa fluor 488 (Invitrogen A21202), donkey anti-rabbit Alexa fluor 546 (Invitrogen, A10040), goat anti-rat Alexa fluor 633 (Invitrogen, A21094), all used 1:1000. When appropriate, F-actin was labeled with Phalloidin Alexa fluor 647 (1:1000, Invitrogen A22287) or Phalloidin rhodamine (Sigma P1951, 1:10,000). Coverslips were mounted in Citifluor MWL4-88-25 (CliniSciences), imaged with Leica SP5 SMD confocal microscope equipped with oil objective 63X HCX Plan Apo CS oil 1.4NA under Metamorph 7.7.6.

### Image quantification and live imaging

Actin organization was evaluated visually after F-actin staining in fixed osteoclasts; a podosome belt was classified abnormal when F-actin staining was fragmented, weak or absent in more than half of the osteoclast periphery, as describe previously (Maurin et al., 2018).

For live imaging, osteoclasts were differentiated from WT and Tubb6 KO RAW264.7 cells expressing either LifeAct-mCherry to follow actin dynamics or EB1-GFP to follow microtubule dynamics. LifeAct-mCherry was subcloned into pMXs-Puro, retroviral particles were produced as described (Guimbal et al., 2019) and RAW264.7 cells were infected for 24h in 8 μg/ml polybrene (Sigma). Alternatively, RAW264.7 were infected with lentiviral particles obtained from EB1-GFP lentiviral vector (Addgene 118084) for 5 hours in 8 μg/ml polybrene (Sigma).

For microtubule regrowth assays, osteoclasts were treated with 10 μM nocodazole on ice for 2.5 hours to depolymerize microtubules. Medium was then replaced by nocodazole-free medium at 37°C. At desired time points, cells were rinsed in PHEM containing 0.1% saponin, 0.25 nM nocodazole and 0.25 nM paclitaxel, and fixed with methanol at −20°C for 5 minutes. After one-hour saturation in 2% BSA in PBS, cells were processed for immunofluorescence to label microtubules with α tubulin antibody (1:2000) and donkey anti-mouse Alexa488 secondary antibody (1:1000). Images were acquired as described in the Immunofluorescence section above. Aster areas were measured using Segmented Line function in ImageJ 1.53c.

For microtubule dynamics, EB1-GFP-expressing osteoclasts were imaged at 37°C with inverted spinning disk Nikon Ti Andor CSU-X1 equipped with EMCCD iXon897 Andor camera, one image per second for 2 minutes, with oil objective 100x Plan Apo lambda 1.45 NA. EBI-GFP comet mean growth speed, mean length and mean lifetime were determined using u-track (V2.0) software. Comet detection settings were 2 and 4 pixels respectively for low-pass and high-pass Gaussian standard deviation. For tracking the maximum gap to close was set to 4 frames while the minimum length of track segment from first step was set to 10 frames. The remaining settings used for tracking were the u-track default ones. Images were also analyzed in ImageJ 1.53c with MTrackJ function microtubule growth speed, as described (Guimbal et al., 2019).

Endogenous EB1 and EB1-GFP comet length were measured with the Straight-Line function of ImageJ 1.53c.

For podosome belt dynamics, LifeAct-mCherry-expressing osteoclasts were imaged at 37°C with Dragonfly spinning disk microscope equipped with an EMCCD iXon888 Life Andor camera (Andor Technology), 2 images per minute, with 40X plan fluor 1,3 NA DT 0.2 μm objective. The overlap of actin ring position between different time points was determined with the function Measure in ImageJ 1.53c.

For individual podosome dynamics, LifeAct-mCherry-expressing osteoclasts were imaged at 37°C with CSU-X1 spinning disk microscope as above, 1 image per second was acquired for 10 minutes, with 100X plan Apo lambda 1.45 NA objective. Podosome life time was determined by tracking manually on the videos the interval between the appearance and disappearance of individual podosomes.

### Preparation of osteoclast microtubule-associated protein enriched fractions

Osteoclasts were lysed in Tub/MT microtubule stabilizing buffer (80 mM MES, 1 mM EGTA, 4M glycerol, 2 mM MgCl_2_, 1 mM Na_3_VO_4_, pH 6.8) supplemented with 0.1% Triton X-100 and protease inhibitory cocktail for 4 minutes at room temperature. Lysates were centrifuged for 10 minutes at room temperature at 16,000 × g. Pellets were resuspended in Tub/MT supplemented with 20 μM paclitaxel, 0.1 μM GTP, 350 mM NaCl and protease inhibitory cocktail and centrifuged at 21,500 × g for 10 minutes. The resulting supernatant S2, enriched in microtubule binding proteins, was further analyzed by proteomics.

### Microtubule co-sedimentation assay

Human embryonic kidney 293T (HEK293T) cells were transfected with either pRK5 HA-GST (Bompard et al., 2018) or pRK5 HA-GST ARHGAP10, PCR-cloned into pRK5 HA-GST using pEGFP-N3 ARHGAP10 as template, a generous gift from Dr Daisuke Mori (Sekiguchi et al., 2020), using jetPEI (Polyplus transfection) following manufacturer instructions. 24 hours post transfection, cells were harvested, washed twice in ice cold 1X PBS and lysed in 1X BRB80 (80 mM PIPES, 1 mM EGTA, 1 mM MgCl2, pH 6.8) supplemented with 1 mM DTT, protease inhibitor cocktail and 0.1 % IGEPAL CA-630 for 20 min on ice. Cell lysates were clarified at 100,000 × g. for 30 min at 4°C. 100 μg of clarified lysates were supplemented with 2 mM GTP, incubated 5 min on ice followed by 10 min at 37°C. 20 μM paclitaxel was added and the reaction incubated at 37°C for 10 more minutes. Reaction was loaded on a 40 % glycerol cushion made in 1X BRB80 supplemented with 20 μM paclitaxel and centrifuged at 100,000 × g. for 15 min at 30°C. Pellet was rinsed once with 1X BRB80 supplemented with 20 μM paclitaxel and resuspended in the same volume as supernatant. For control experiments, all steps were performed at 4°C in the absence of GTP and Paclitaxel. Samples were then analyzed by immunoblot.

### Proteomics

After reduction (DTT 1M, 30 min at 60°C) an alkylation (IAA 0.5M, 30 min RT) microtubule-associated protein enriched fractions were digested using trypsin (Gold, Promega, 1 μg / sample, overnight at 30°C). Peptide clean-up was done using OMIX C18 (Agilent) according to manufacturer protocol. For LC MSMS anlysis, samples were loaded onto a 25 cm reversed-phase column (75 mm inner diameter; Acclaim PepMap 100 C18; Thermo Fisher Scientific) and separated with an UltiMate 3000 RSLC system (Thermo Fisher Scientific) coupled to a QExactive HFX system (Thermo Fisher Scientific). Tandem mass spectrometry analyses were performed in a data-dependent mode. Full scans (350–1,500 m/z) were acquired in the Orbitrap mass analyser with a resolution of 120,000 at 200 m/z. For MS scans, 3e6 ions were accumulated within a maximum injection time of 60 ms. The 20 most intense ions with charge states ≥2 were sequentially isolated (1e5) with a maximum injection time of 50 ms and fragmented by higher-energy collisional dissociation (normalized collision energy of 28) and detected in the Orbitrap analyzer at a resolution of 30,000. Raw spectra were processed with MaxQuant v 1.6.10.43 (Cox and Mann, 2008) using standard parameters with LFQ and match between runs option. Spectra were matched against the UniProt reference proteome of Mus musculus (UP000000589, release 2020_04; http://www.uniprot.org) and 250 frequently observed contaminants, as well as reversed sequences of all entries. The maximum false discovery rate for peptides and proteins was set to 0.01. Representative protein ID in each protein group was automatically selected using the in-house developed Leading tool v 3.4 (Raynaud et al., 2018). Statistical analysis (t-test) were done using Perseus v 1.6.10.43 (Tyanova et al., 2016). Briefly, after reverse and contaminants entries removal, LFQ data were log2 transformed. Imputation was performed using “Replace missing values from normal distribution” with standard parameters. Difference between the 2 groups was then assessed using a standard t-test.

GO term enrichment analyses were performed using Panther 16.0 software (pantherdb.org/, GO Ontology database DOI: 10.5281/zenodo.5080993 Released 2021-07-02). UniProtIDs were employed as input type IDs and the total list of Mus musculus genes was used as reference. GO cellular component, molecular function and cellular component terms with P<0.05 with Fisher’s Exact were considered significantly enriched.

### Statistical analyses

Statistical significance was assessed either with parametric statistical test after normality assessment or with non-parametric tests. All analyses were done with GraphPad Prism 9.2.0 (GraphPad Software, Inc.), with p < 0.05 considered statistically significant.

## RESULTS

### TUBB6 impacts microtubule dynamics in osteoclasts

To understand how TUBB6 controls actin organization and bone resorption, we generated RAW264.7 cells KO for Tubb6 by CRISPR/Cas9 from which we derived osteoclasts (Figure 1A–B and supplementary Figure 1A). Similar to what we described in primary osteoclasts treated with Tubb6 siRNAs (Guérit et al., 2020), we found that osteoclasts derived from RAW264.7 Tubb6 KO cells presented several defects when plated on glass: increase of abnormal podosome belts and podosome rings, which are immature podosome belts (Figure 1C–D and Supplementary Figure 1B). Consistently, Tubb6 KO led to osteoclasts with less sealing zones on mineralized substrates and more rings, which are immature sealing zones (Figure 1E–F and supplementary Figure 1C), as well as reduced resorption activity (Figure 1G). The induction of the expression of osteoclastic markers Src and CtsK was not affected during Tubb6 KO osteoclast differentiation (Supplementary Figure 1D–E), global expression of β tubulin (Figure 1A) and the expression of β tubulin isotype TUBB5 were also unchanged (Supplementary Figure 1F). Altogether, this shows that RAW264.7 osteoclasts KO for Tubb6 recapitulate the features of primary osteoclasts treated with Tubb6 siRNAs.

**Figure 1:**
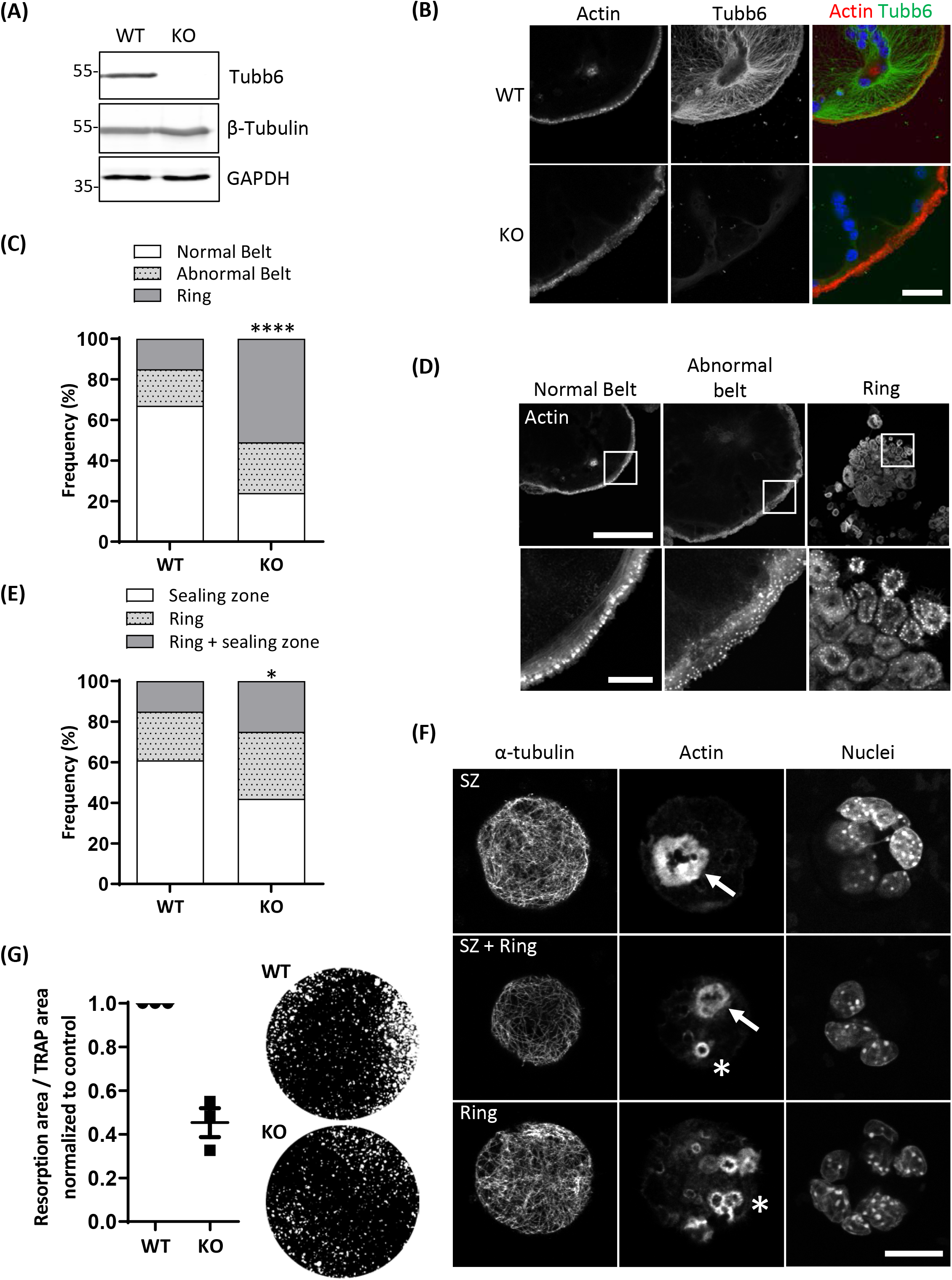
Tubb6 gene knockout alters osteoclast actin cytoskeleton and activity. Representative immunoblot analyses of lysates from wild type (WT) and Tubb6 KO (KO) osteoclasts. GAPDH control shows that total β-tubulin levels are not affected in Tubb6 KO osteoclasts. (B) Representative maximum intensity projection (MIP) confocal images of WT and KO osteoclasts seeded on glass and stained for endogenous F-actin and TUBB6. Scale bar: 40 μm. (C) Bar graph showing the frequency of WT and KO osteoclasts seeded on glass and presenting podosome rings, normal or abnormal podosome belt, counting more than 200 WT and KO osteoclasts in 4 independent experiments, χ^2^ contingency test, ****p<0.0001. (D) Representative MIP images illustrating the different categories of actin organization considered in C. Scale bars are respectively 100 μm and 20 μm for top and bottom images. (E) Bar graph showing the frequency of WT and KO osteoclasts seeded on mineral matrix and presenting either sealing zones (SZ), or rings or both, counting more than 150 WT and KO osteoclasts in 3 independent experiments, χ^2^ contingency test, *p<0.05. (F) Representative MIP images illustrating the different categories of actin organization considered in E, with SZ and rings respectively indicated with arrows and stars respectively. Scale bar: 20 μm. (G) Graph showing the average and SEM specific resorption activity of KO osteoclasts normalized to the activity of WT osteoclasts in three independent experiments, 4 wells per experiments. Representative image of von Kossa staining of the resorption wells are presented on the right.

To assess the impact of TUBB6 on osteoclast microtubule dynamics, we first performed microtubule regrowth assay consisting in studying microtubule polymerization after cold and nocodazole treatment followed by nocodazole washout. We found that microtubule regrowth was slower in Tubb6 KO osteoclasts (Figure 2A–B). To confirm this, we used Tubb6 siRNAs to lower TUBB6 expression (Guérit et al., 2020). Again, we found that Tubb6 siRNAs reduced the speed of microtubule regrowth (Supplementary Figure 2). To get additional information regarding this defect, we performed live imaging of osteoclasts expressing the microtubule plus tip-binding protein EB1 fused to GFP (Figure 3A). Using time laps microscopy and u-track software, we tracked EB1-GFP comets on individual microtubule (Figure 3B and Supplementary Videos 1 & 2) and measured microtubule dynamic parameters. Thereby, we found a significant decrease of microtubule growth speed in Tubb6 KO osteoclasts from a mean value of 16 μm/min in the wild type down to 12.9 μm/min in the KO (Figure 3C and Supplementary Figure 3A). Similar results were found by manual tracking using the MTrack plugin in ImageJ (Supplementary Figure 3B–D). Conversely, we found an increase in microtubule growth time from 18.1 seconds in the wild type to 20.3 in the KO, but no significant change in microtubule growth length (Figure 3C and Supplementary Figure 3A). We also observed that the level of tubulin acetylation, a marker of microtubule stability, was higher in Tubb6 KO osteoclasts (Figure 3D–E), potentially consistent with longer microtubule growth time. Finally, we measured the size of endogenous EB1 staining at microtubule ends, as indicative of the length of the microtubule GTP cap that is known to be proportional to growth speed. We found that the length of EB1 staining was reduced form 1.393 μm in wild type down to 1.089 μm in Tubb6 KO osteoclasts (Figure 3F), consistent with their lower microtubule growth speed (Vemu et al., 2017). The length of the EB1-GFP staining at the end of microtubules was similarly reduced in Tubb6 KO osteoclasts (Supplementary Figure 3E–G).

**Figure 2:**
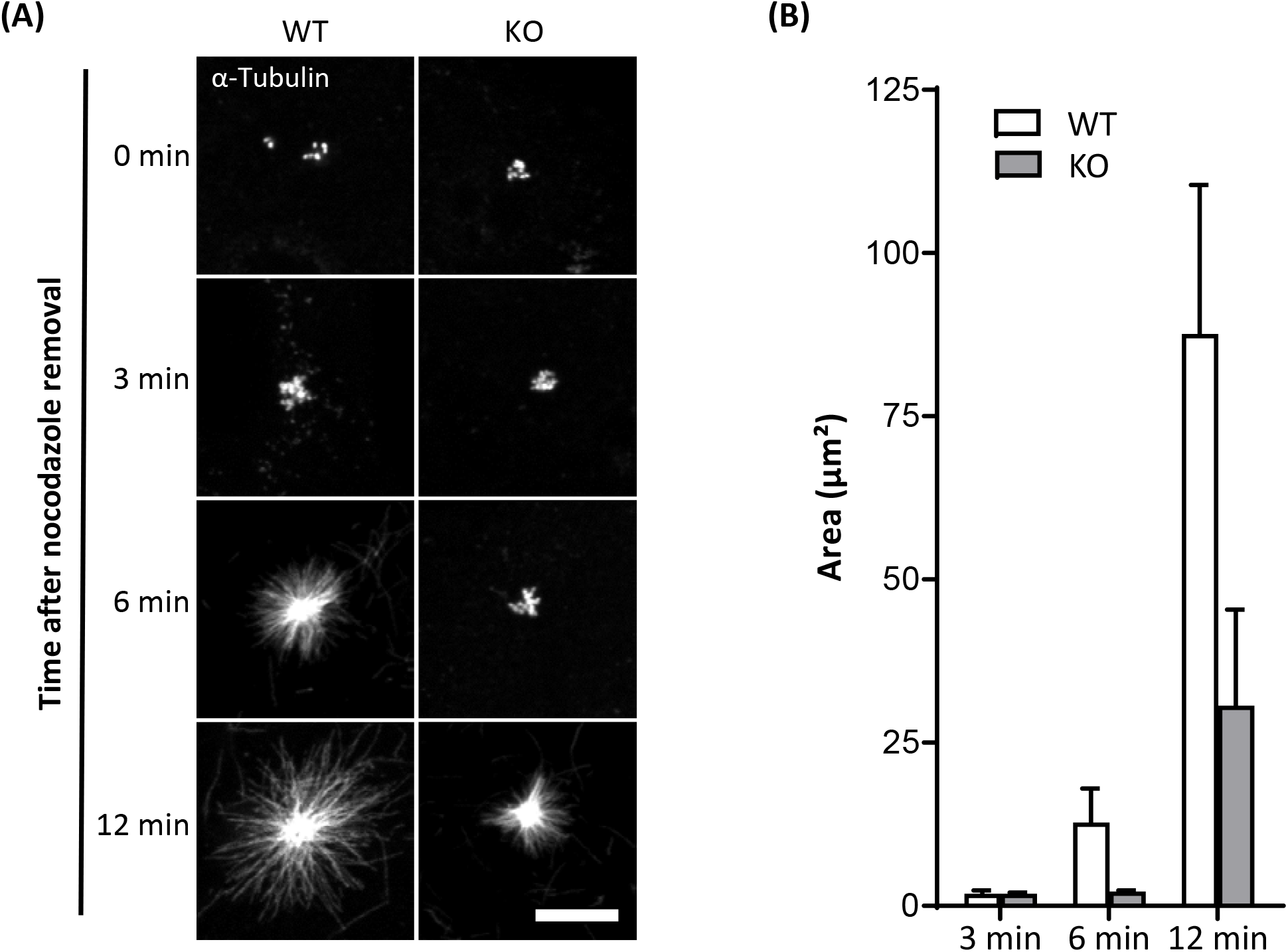
Tubb6 depletion affects MT polymerization. (A) Representative MIP images of WT and KO osteoclasts stained for α-tubulin, after release from nocodazole and cold treatment for indicated times. Scale bar: 10 μm. (B) Bar graph showing mean and SEM of α-tubulin aster area in WT and KO osteoclasts treated as in (A), 80 WT and KO asters in 4 independent experiments.

**Figure 3:**
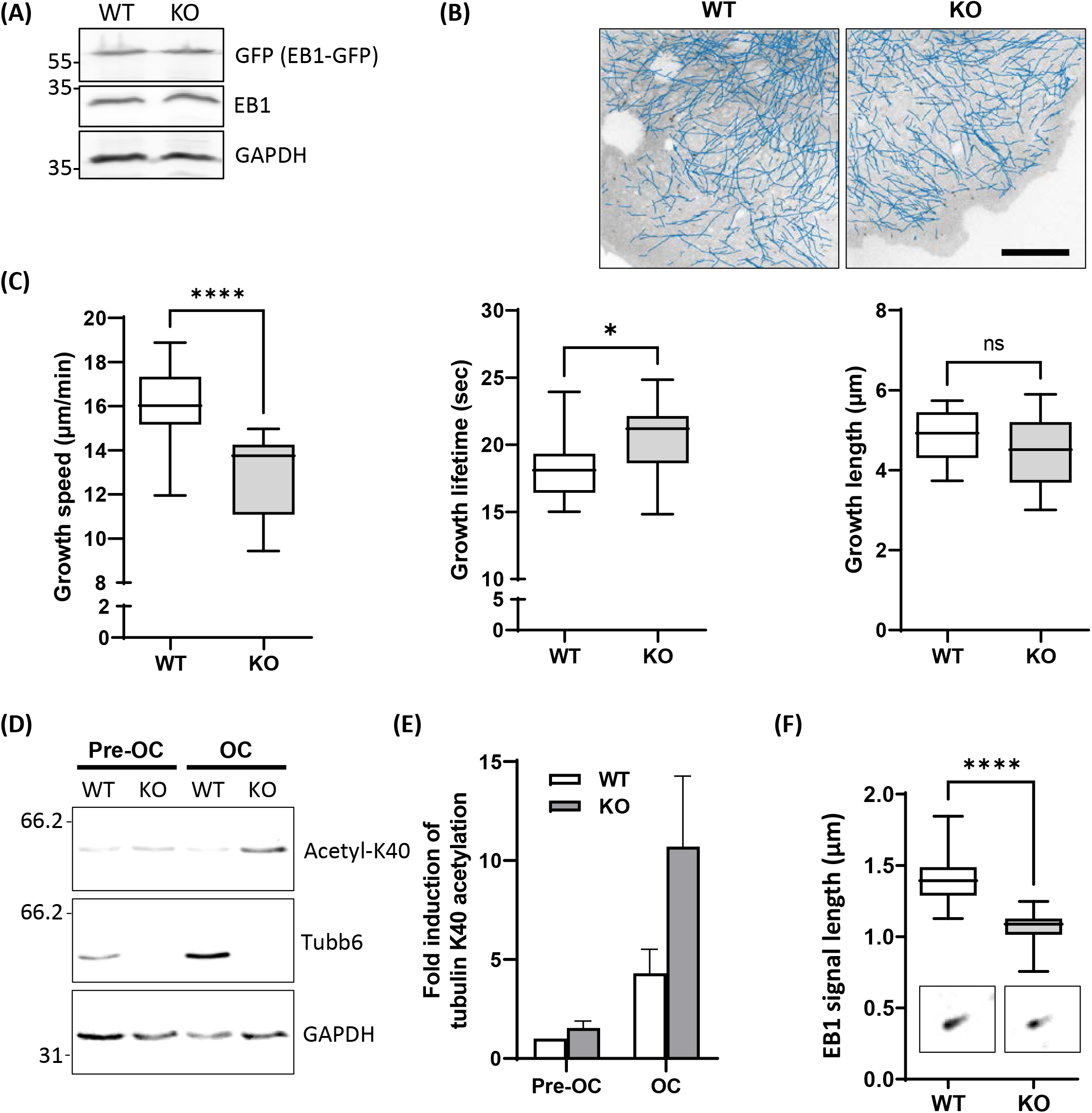
Tubb6 regulates MT dynamics. (A) Representative immunoblot analysis of protein lysates from WT and KO osteoclasts expressing EB1-GFP (GFP); note that EB1-GFP and endogenous EB1 expressions are similar in WT and KO osteoclasts. (B) Blue lines show representative EB1-GFP comet tracking using u-track software in WT and KO osteoclasts from confocal images over a period of 2 minutes with 1 second increment between frames (see material and methods for detailed settings). Scale bar: 20 μm. (C) Minimum to maximum boxplots with median and interquartile range (25-75%) showing the distribution of growth speed (left), lifetime (middle) and length (right) of EB1-GFP tracked comets per single WT and KO osteoclast, 18 WT and 14 KO osteoclasts from 3 independent experiments; Mann Whitney test: ****p<0.0001 and *p<0.05. For details see table in Supplementary figure 3A. Scale bar: 10 μm. (D) Representative immunoblot analysis of protein lysates from WT or KO RAW264.7 cells (Pre-OC) and derived osteoclasts (OC), showing TUBB6, acetylated K40 α-tubulin (acetyl-K40) and GAPDH expression. (E) Bar graph shows mean and SEM levels of acetylated K40 α-tubulin in WT and KO pre-OC and OC, normalized to 1 in WT pre-osteoclasts, from 4 independent experiments. (F) Minimum to maximum boxplot showing endogenous EB1 cap length in WT and KO osteoclasts, measuring 25 comets in each osteoclast to determine the average cap length per osteoclast, in 27 WT and 27 KO osteoclasts from 3 different experiments; Mann Whitney test: ****p<0.0001.

These results show that TUBB6 has an impact on the parameters of microtubule dynamics, increasing microtubule growth speed while decreasing microtubule lifetime.

### TUBB6 impacts on actin dynamics in osteoclasts

Actin cytoskeleton and microtubules are intimately linked and Tubb6 KO osteoclasts show abnormal actin cytoskeleton organization. Thus, we further examined the impact of TUBB6 on actin dynamics, by performing live imaging of osteoclasts expressing LifeAct-mCherry to visualize actin cytoskeleton (Figure 4A, Supplementary Figure 4 and Supplementary Videos 3 & 4). Major differences in podosome belt dynamics were observed and we thus assessed the stability of the podosome belt by comparing the position of LifeAct-mCherry labeling between time points. In control osteoclasts, the median overlap of LifeAct-mCherry staining between time 0 and time 1 = 3 minutes was 69.56%, whereas this overlap was significantly reduced to 55.79% in Tubb6 KO osteoclasts (Figure 4A–B). Similar observations were made at time 2 = 6.5 minutes (Supplementary Figure 4). Finally, we measured the life span of isolated podosomes and found that Tubb6 KO increased to 3.31 minutes, instead of 2.5 minutes in the wild type (Figure 4C and Supplementary Videos 5 & 6).

**Figure 4:**
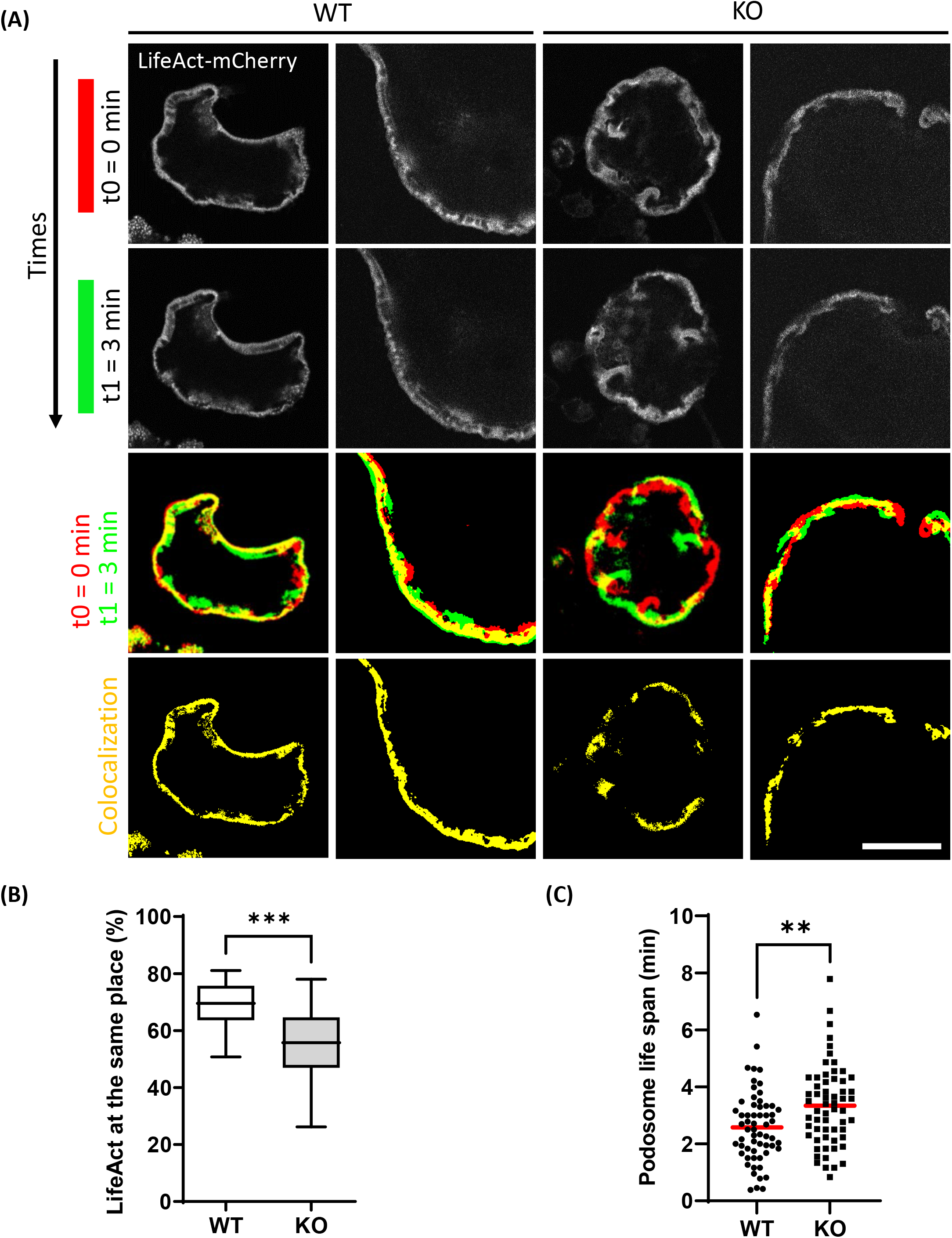
Tubb6 depletion affects podosome belt dynamics. (A) Representative live confocal images of osteoclasts derived from WT or Tubb6 KO osteoclasts expressing LifeAct-mCherry and sitting on glass, showing the localization of LifeAct-mCherry signal at t0 = 0 minute (red) and t1 = 3 minutes (green), with the overlapping areas in yellow. Scale bar: 30 μm. (B) Minimum to maximum boxplot showing the percentage of LifeAct-mCherry signal at t0 that persists at the same position at t3 in WT or Tubb6 KO osteoclasts; in a total of 22 WT and 21 KO osteoclasts from 4 different experiments. Mann Whitney test: ***p< 0.0002. See also Supplementary Figure 4 and supplementary videos 3 and 4. (C) Dot plot showing the life span with mean of individual podosomes in WT or Tubb6 KO osteoclasts. Total of 60 podosomes in 6 different wild-type and KO osteoclasts, from 3 independent experiments. Student’s t-test: **p<0.002.

This suggests that, on top of microtubules, TUBB6 also impacts on actin dynamics in osteoclasts: TUBB6 tends to reduce podosome life span whereas it increases the stability of the podosome belt. Altogether, the data show that TUBB6, which is required for correct podosome patterning and bone resorption, influences both actin and microtubule dynamics in osteoclasts.

### TUBB6 moderately affects protein binding to osteoclast microtubules

To understand the impact of TUBB6 on osteoclast cytoskeleton, we examined whether Tubb6 KO could affect protein binding to osteoclast microtubules. For this, we thought to prepare osteoclast fractions enriched in microtubule-associated proteins (Figure 5A). Osteoclasts lysates were prepared at room temperature, in a paclitaxel-free microtubule-preserving hypotonic buffer. Lysates were centrifuged to separate the supernatant (S1) from the pellet. The pellet was then washed with the same buffer supplemented with 350 mM NaCl, GTP and paclitaxel, to detach microtubule-bound proteins (S2) from microtubules while preserving the latter, which were finally pelleted (P) (Figure 5A). Immunoblot analysis of the different fractions showed the expected marker distribution: total tubulin equally distributed in S1 and P whereas acetylated tubulin fell predominantly in P, which contains microtubules; the microtubule-associated protein CLASP1 was found in S2 whereas vinculin was predominantly in S1 (Figure 5B). Of note, all fractions contained similar amounts of actin, indicative of partial F-actin depolymerization in 350 mM NaCl (Figure 5B). Finally, histone H3 was found in the pellet, in agreement with the presence of nuclei in this fraction; small amounts of histone H3 were also found in S2, likely due to nuclear protein leaking out of the nucleus upon high salt incubation after hypotonic lysis of osteoclasts (Figure 5B).

**Figure 5:**
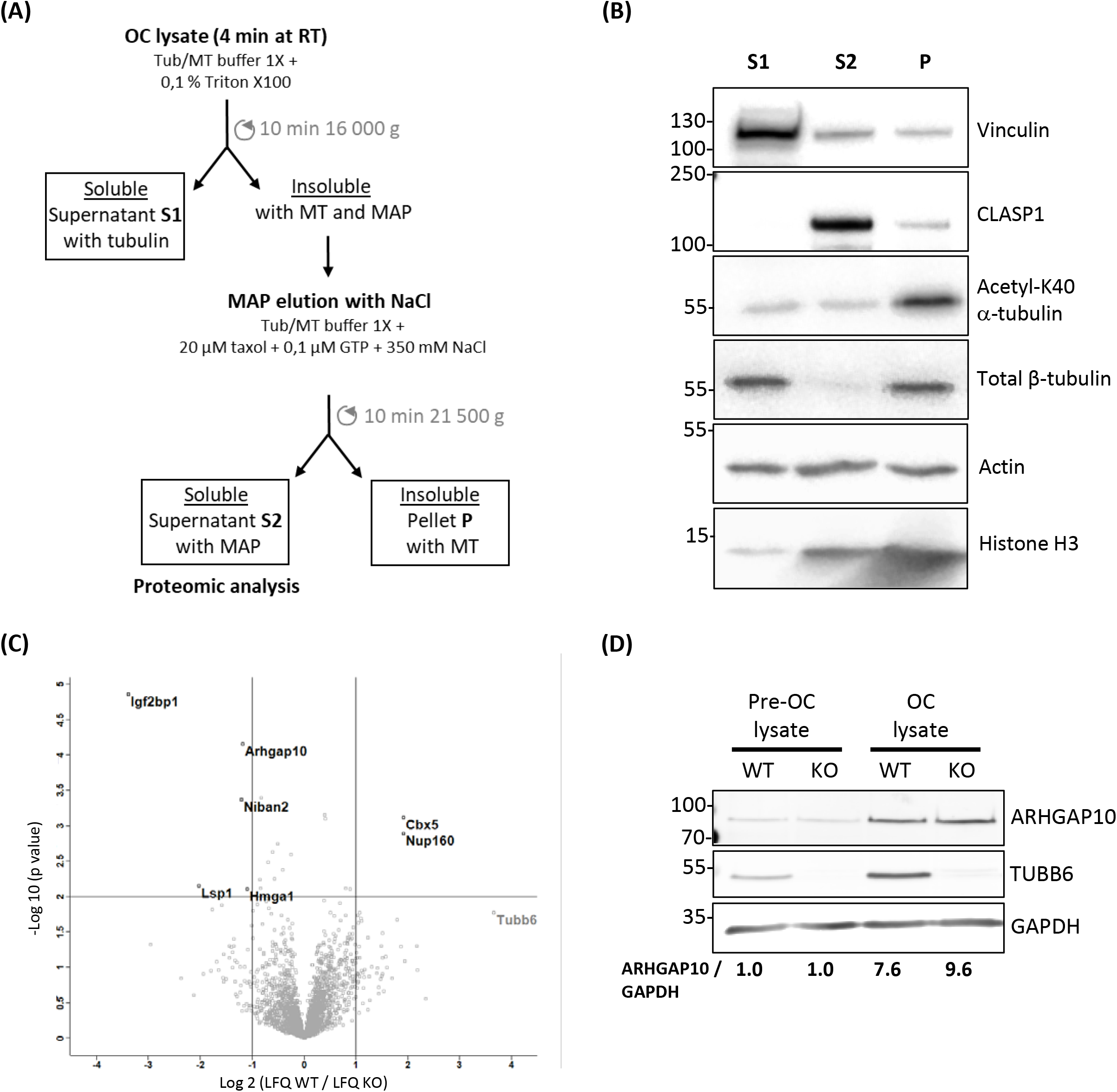
Characterization of the microtubule-associated protein fraction of WT and Tubb6 KO osteoclasts. (A) Schematic representation of the osteoclast fractionation strategy in microtubule-stabilizing Tub/MT buffer to obtain soluble proteins (S1), a fraction enriched in microtubule-binding protein (S2) and an insoluble fraction containing microtubules (P). (B) Representative immunoblot analysis of selected protein distribution between the different fractions. (C) Volcano plot of Label Free Quantitation (LFQ) results from 4 independent mass spectrometry experiments, showing proteins that are significantly more abundant in S2 from KO (top left) and WT (top right) osteoclast, with vertical lines indicating 2-fold change cut off and horizontal line p=0.01 cut off, with x axis in log2(LFQ WT / LFQ KO) and y axis in −log10(p-value). (D) Representative immunoblot analysis of ARHGAP10 expression in from WT or KO RAW264.7 cells (Pre-OC) and derived osteoclasts (OC) with indicated antibodies.

We performed label-free proteomics to compare protein content in the S2 fractions of wild type and Tubb6 KO osteoclasts, in 4 independent experiments. We identified 2799 protein groups, considering 2 peptides for adequate protein identification, among which 2778 corresponded unambiguously to a single Uniprot identifier and to a unique protein (Table S1). To examine whether S2 fraction was enriched in cytoskeleton regulatory proteins, we used Panther 16.0 to perform gene ontology analyses of the 2773 proteins (5 Uniprot identifiers could not be mapped in Panther), as compared to the total mouse genome (Table S2). Using actin, tubulin, microtubule and cytoskeleton as keywords, we found that all Gene Ontology (GO) terms associated with the cytoskeleton in molecular functions, cellular components and biological process showed a significant enrichment in the S2 fraction (Table S2). Cytoskeleton-related proteins associated with these GO terms encompassed 601 of the 2773 proteins we analyzed in Panther (Table S3). Thus, S2 contains a great proportion of proteins related to microtubules and actin cytoskeleton.

Applying a 2-fold change cutoff with a p-value below 10^−2^, we found only 7 proteins showing significant changes in the S2 of Tubb6 KO osteoclasts as compared to WT (Figure 5C and Table 1). Among these, only 2 were part of the 601 proteins associated with cytoskeleton-related GO terms: the podosome-associated protein LSP1 (lymphocyte specific 1) and the Rho-family GTPase regulating protein ARHGAP10 (also known as GRAF2 or PS-GAP) (Figure 5C and Table 1).

**Table 1:**
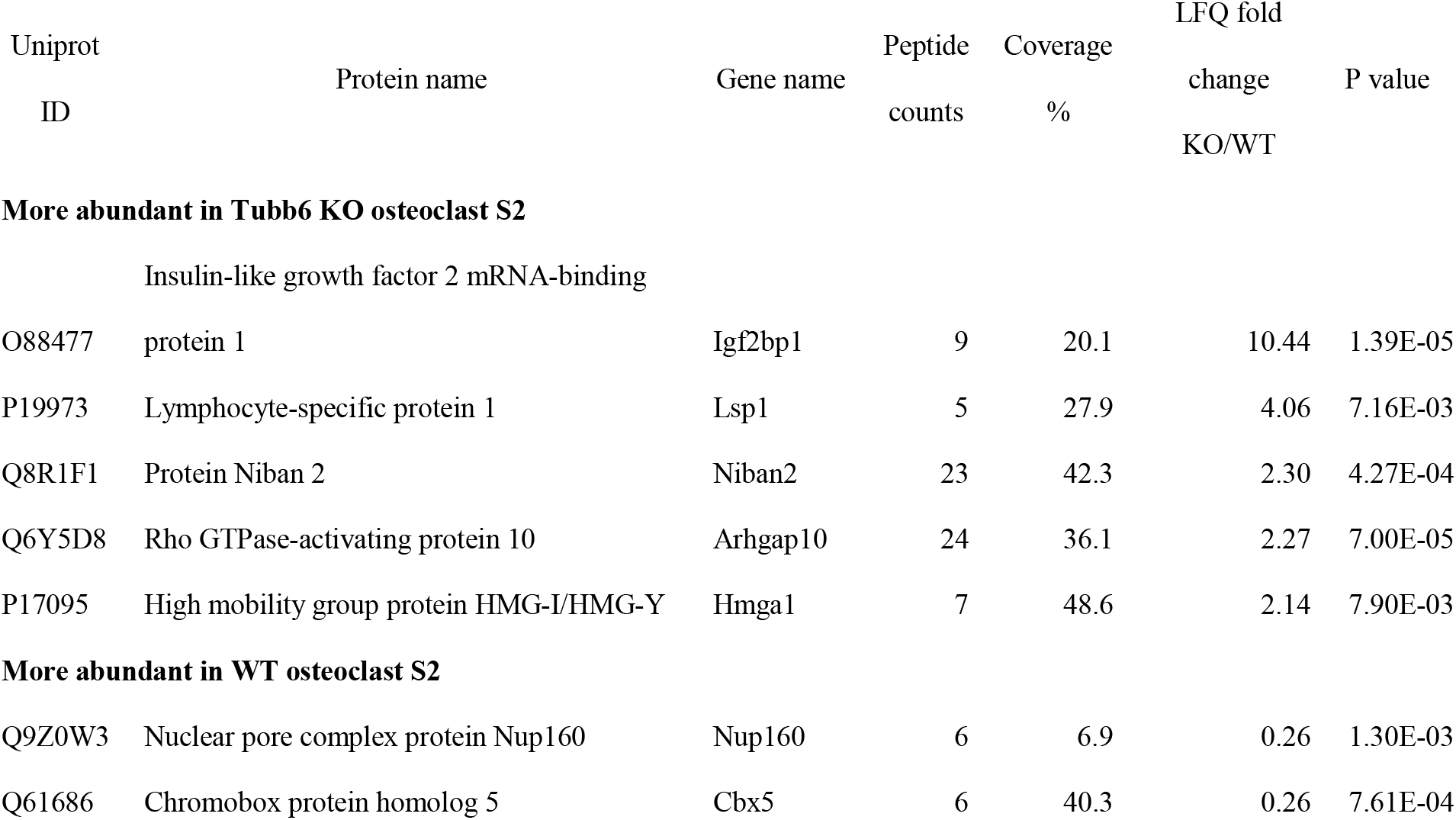
Protein changes in fractions S2.

These results show that TUBB6 has little effect on the content of the osteoclast S2 fraction, which is enriched in microtubule-associated protein. In particular, among the 601 proteins present in S2 and associated with a cytoskeleton-related GO term, only LSP1 and ARHGAP10 showed significant changes. This suggests that TUBB6 can affect the interaction of LSP1 and ARHGAP10 with F-actin and/or microtubules.

### ARHGAP10 associates with both actin and microtubules in osteoclasts

ARHGAP10 was shown to be a negative regulator of the activity of the GTPases CDC42, in a PYK2-kinase dependent fashion (Koeppel et al., 2004; Ren et al., 2001); ARHGAP10 can also regulate negatively the activity of GTPase RHOA (Ren et al., 2001). It was reported in RAW264.7-derived osteoclasts that Arhgap10 siRNAs led to smaller sealing zones, without affecting the proportion of osteoclasts with a sealing zone (Steenblock et al., 2014). Interestingly, we also observed smaller sealing zones upon Tubb6 KO (Figure 1 and supplementary Figure 1), as well as with Tubb6 siRNAs in primary mouse and human osteoclasts (Guérit et al., 2020). We found that ARHGAP10 expression was strongly induced during both WT and Tubb6 KO osteoclasts (Figure 5D). Endogenous ARHGAP10 colocalized with the podosome belt in RAW264.7-derived and primary osteoclasts, as well as with the sealing zone of osteoclasts plated on ACC (Figure 6A–B). Interestingly, we also observed endogenous ARHGAP10 colocalization with osteoclast microtubules (Figure 6C–D). To confirm that ARHGAP10 can interact with microtubules, we overexpressed ARHGAP10 fused to a HA-GST tag in HEK293T cells, which do not express endogenous ARHGAP10 (Koeppel et al., 2004), or HA-GST as a control. After lysis in a buffer compatible with microtubule polymerization and clarification by high-speed centrifugation, lysates were either incubated at 4°C or 37°C in the presence of GTP and paclitaxel to induce microtubule polymerization. Finally, the fraction of HA-GST fused proteins co-sedimenting with microtubules were analyzed by immunoblot after a final high-speed centrifugation. Tubulin was found only in the 37°C-pellet, indicating that no microtubule polymerized at 4°C. As expected, HA-GST was only retrieved in supernatants regardless the temperatures (Figure 6E). Interestingly, at 37°C, HA-GST ARHGAP10 almost exclusively co-sedimented with microtubules whereas it was retained in the supernatant when the experiment was performed at 4°C to prevent microtubule polymerization (Figure 6F). Of note, very little actin was retrieved in the osteoclast lysate and no actin could be detected in the pellets that contained ARHGAP10 (Figure 6F). These data show that ARHGAP10 is indeed a microtubule-associated protein.

**Figure 6:**
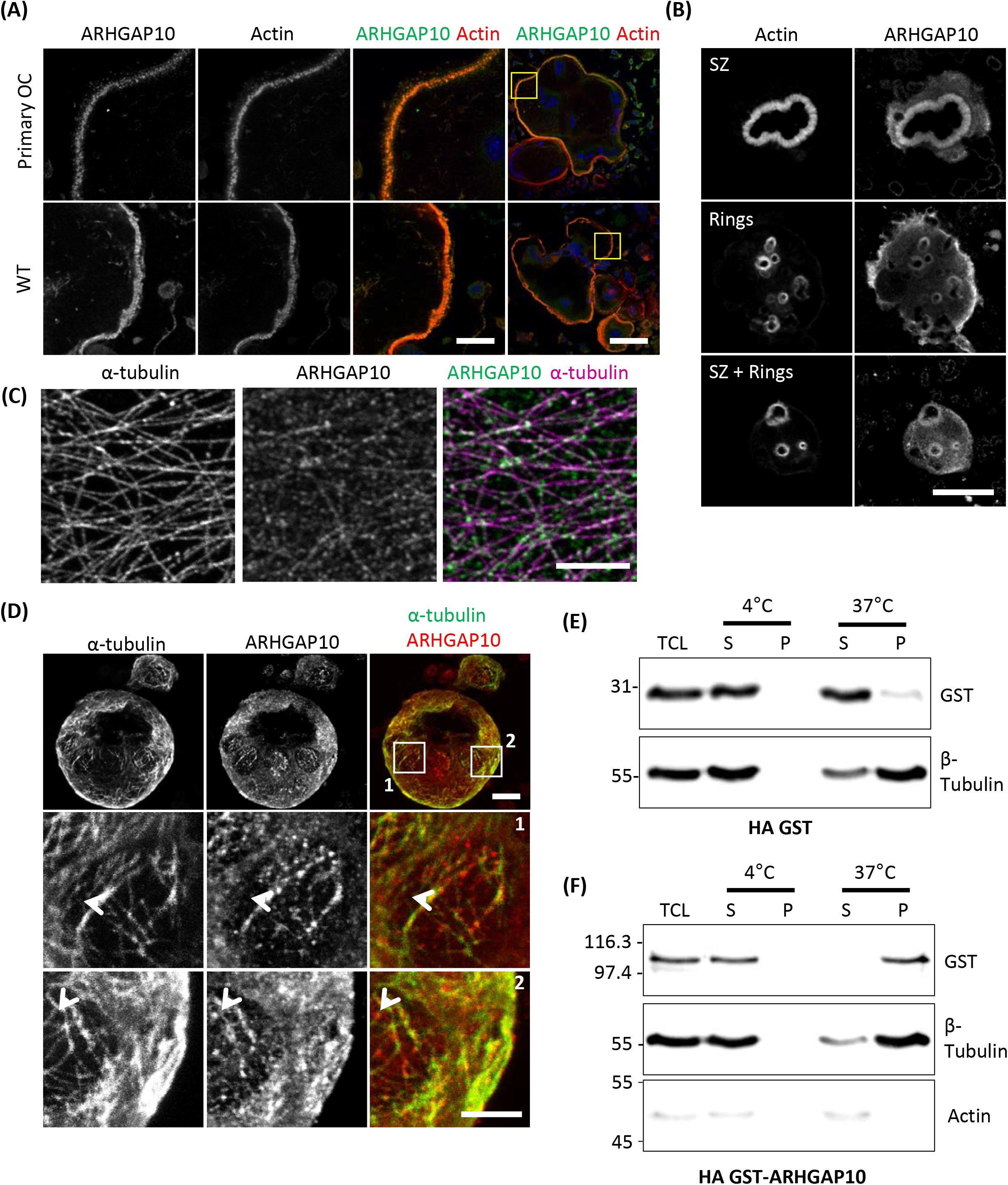
ARHGAP10 is associated with both actin and microtubules. (A) Representative MIP images of RAW264.7 derived or primary (primary OC) osteoclasts seeded on glass and stained for ARHGAP10, actin and DNA. Left images show enlargement of boxed areas in the right images. Scale bars: 20 μm for right images and 100 μm for other images. (B) Representative MIP images showing ARHGAP10 and actin localization in WT and Tubb6 KO osteoclasts sitting on ACC. Scale bar: 20 μm. (C) Representative single plan confocal image of a WT osteoclast stained for ARHGAP10 (green) and α-tubulin (pink). Scale bar: 5 μm. (D) Representative MIP images of a WT osteoclast sitting on ACC and stained for ARHGAP10 and α-tubulin. Boxed areas in top images are enlarged below, to show ARHGAP10 colocalization with microtubules (arrowheads). Scale bar: 10 μm in top images and 5 μm in other images. (E-F) Immunoblot analysis of lysates from HEK293T cells expressing HA-GST (E) or HA-GST-ARHGAP10 (F) and subjected to microtubule pelleting assays at 4°C or 37°C. ARHGAP10 pellets with microtubules, which are in the 37°C pellet.

Altogether, our data suggest that TUBB6 can control the amount of ARHGAP10 associated with microtubules in osteoclasts. This may contribute to control the size of sealing zones, which was shown to involve both TUBB6 (Guérit et al., 2020) and ARHGAP10 (Steenblock et al., 2014).

## DISCUSSION

The cytoskeleton plays a key role in bone resorption by osteoclasts. F-actin organizes as a belt of podosomes required for adhesion and as backbone of the ruffled border, which is the actual bone resorption apparatus. Microtubules are also instrumental for podosome organization in osteoclasts and bone resorption but the coordination mechanisms between the actin and tubulin cytoskeletons remain poorly characterized.

We identified previously TUBB6 as a β-tubulin isotype essential in osteoclasts for podosome belt integrity and for resorption (Guérit et al., 2020). To study the implications of TUBB6 in cytoskeleton dynamic parameters, we developed a model of osteoclasts KO for Tubb6 using the RAW264.7 pre-osteoclast cell line, which recapitulates the phenotype we reported recently for primary osteoclasts treated with Tubb6 siRNAs (Guérit et al., 2020). On top of the expected abnormal actin organization and reduced resorption activity, we found that the lack of TUBB6 also effected microtubule dynamics in osteoclasts: the speed of microtubule repolymerization after nocodazole treatment was reduced as well as individual microtubule growth speed, whereas their lifetime was increased, accompanied by an increase in tubulin acetylation. This suggests that TUBB6 has a microtubule-destabilizing effect in osteoclasts. Interestingly, overexpression of TUBB6 was reported previously to disrupt microtubule organization in various cycling cell types (Bhattacharya and Cabral, 2004) and to induce paclitaxel resistance, with serine 239 being essential for TUBB6 microtubule-destabilizing activity (Bhattacharya and Cabral, 2009). Serine 239 is located within the core of β-tubulin and it is conserved in TUBB1 and TUBB3. Interestingly, similarly to TUBB6, high expression of TUBB3 is suspected to participate in paclitaxel resistance (Kato et al., 2018). Furthermore, TUBB3 has been shown to significantly decrease microtubule growth rates (Pamula et al., 2016; Vemu et al., 2016), similar to what we found here for TUBB6. Furthermore, this intrinsic property on microtubule dynamics does not require residues located within the carboxy-terminal tail of TUBB3 leaving the possibility of serine 239 involvement (Pamula et al., 2016). Based on these data, it is very likely that TUBB6 intrinsically alters the dynamics of microtubules, similar to TUBB3.

We noticed an increase of K40-acetylation of α-tubulin upon KO of Tubb6 in osteoclasts, as also observed in CHO cells expressing Tubb6 shRNAs (Bhattacharya et al., 2011). K40-acetylation of α-tubulin is a well described marker of stable MT (Janke and Magiera, 2020). The increase of K40-acetylation we observed in Tubb6 KO osteoclasts may thus be a consequence of the increase of MT lifetime that we measured. K40-acetylation has been shown to play a crucial role in podosome belt establishment and/or stability in osteoclasts (Destaing et al., 2005). Lower K40-acetylation levels were also associated with various genetic defects leading to the alteration of podosome patterning in osteoclasts, such as the KO of Pyk2, myosin IXB or cofilin 1 (Gil-Henn et al., 2007; McMichael et al., 2014; Zalli et al., 2016). In the case of Tubb6 gene disruption, we found conversely an increase of K40-acetylation associated with an abnormal organization of podosomes. Microtubule acetylation was shown to enhance their flexibility (Portran et al., 2017) and actually microtubules are heavily buckled in the podosome belt area of primary osteoclasts treated with Tubb6 siRNAs as compared to control (Guérit et al., 2020). Thus, an optimal level of microtubule acetylation appears essential for osteoclast function, potentially to provide microtubules with appropriate flexibility and mechanical resistance (Portran et al., 2017; Xu et al., 2017). Such optimal balance is known for other PTMs in general and for microtubule PTMs in particular. For instance, increasing and decreasing the level of microtubule glutamylation equally impair ciliary function in the nematode (O’Hagan et al., 2017). K40-acetylation could also participate in the changes in podosome dynamics we observe in Tubb6 KO osteoclasts. Indeed, dynamic microtubules were shown to regulate podosome fate in macrophages: podosome contact by plus-end MT mostly induces their dissolution or fission (Kopp et al., 2006) and increasing K40-acetylation of microtubules diminishes microtubule plus ends targeting of podosomes (Bhuwania et al., 2014). In Tubb6 KO osteoclasts, we observe increased podosome life span correlated with increased microtubule acetylation, which may result from a reduction in contacts between microtubules and podosomes.

Apart from directly impacting on the dynamics of microtubules, tubulin isotypes influence the recruitment of proteins on microtubules, which can also modulate their dynamics as well as cargo transport (Janke and Magiera, 2020). Our proteomic analyses of osteoclast S2 fractions, enriched in microtubule-associated proteins, did not reveal major changes whether osteoclasts expressed TUBB6 or not. Still, we found two cytoskeleton-related proteins: ARHAP10 and LSP1, which were increased in S2 in the absence of TUBB6. Similar to what was reported with Arhgap10 siRNAs (Steenblock et al., 2014), we found that the osteoclast KO of Tubb6 have small sealing zones that often remain immature. ARHGAP10 is a GAP that negatively regulates the activity of CDC42 (Ren et al., 2001). It was shown in osteoclasts to participate in a complex with the CDC42 exchange factor FGD6, all three proteins localizing at the sealing zone (Steenblock et al., 2014). Our data show that ARHGAP10 also colocalizes with osteoclast microtubules and that ARHGAP10 associates very efficiently with microtubules in an actin-independent fashion, suggesting that ARHGAP10 can associate with both microtubules and actin cytoskeleton in osteoclasts. CDC42 is essential for the correct dynamics of actin rings in osteoclasts and for bone resorption *in vitro* and *in vivo* in the mouse; conversely excessive activity of CDC42 causes osteoclast hyperactivity and osteoporosis (Ito et al., 2010). Thus, by controlling ARHGAP10 association with the cytoskeleton, TUBB6 could contribute to the fine tuning of local CDC42 activity to ensure an optimal level of resorption. LSP1 is an actin-regulating protein, which was recently shown to associate with the cap of podosomes in macrophages (Cervero et al., 2018). LSP1 regulates macrophage podosome properties: the levels of LSP1 control the frequency and amplitude of podosome oscillations, and interfering with LSP1 expression reduces podosome life span and the protruding forces they exert on the extracellular matrix (Cervero et al., 2018). Moreover, lack of LSP1 was recently shown to affect the microtubule network in macrophages (Schäringer et al., 2021). Here we found that in the absence of TUBB6, podosome have longer life-span and we found increased amounts of LSP1 in the S2 fraction of Tubb6 KO osteoclasts, suggesting that LSP1 may participate in the control of actin-dynamics in osteoclasts in a TUBB6-dependent fashion. The function of LSP1 in osteoclasts would deserve further investigations.

To conclude, our data highlight the crucial role of microtubule dynamics in podosome belt integrity in general and the involvement of TUBB6 in particular. We provide new insights into the control of microtubule dynamics in osteoclasts and position β-tubulin isotype TUBB6 as a key regulator of the crosstalk between actin and microtubules in osteoclast. Our results uncover novel regulatory mechanisms of osteoclast cytoskeleton and contribute to the better knowledge of the molecular processes controlling bone resorption. Our data could be valuable also in the context of osteolytic bone diseases and pave the way for novel solutions against osteoporosis by targeting osteoclast cytoskeleton.

## AUTOR CONTRIBUTIONS

Conceptualization: A.B., G.B; Methodology: A.B., G.B, J.M, A.M., D.G., J.C., S.U.; Validation: G.B, J.M, A.M., D.G., J.C., S.U.; Formal analyses: A.B., G.B, J.M, A.M., D.G., S.U; Investigation: G.B, J.M, A.M., D.G., S.U; Writing - original draft: A.B., G.B, J.M., S.U.; Supervision: A.B., G.B.; Project administration: A.B.; Funding acquisition: A.B.

## ACKNOWLEDGEMENTS

We acknowledge the imaging facility Montpellier Ressources Imagerie (MRI), a member of the national infrastructure France-BioImaging supported by the French National Research Agency (ANR-10-INBS-04, ‘Investments for the future’). Mass spectrometry experiments were carried out using the facilities of the Montpellier Proteomics Platform (PPM, BioCampus Montpellier). We are very grateful to Ariane Abrieu, Leslie Bancel-Vallée, Daniel Bouvard, Juliette van Dijk, Nathalie Morin, Didier Portran and Amélie Viel (CRBM Montpellier, France) to for technical assistance, helpful advice and discussions. We also thank Dr Daisuke Mori (Nagoya University, Japan) for the generous gift of ARHGAP10 expression vector.

## SUPPLEMENTARY FIGURE AND VIDEO LEGENDS

**Supplementary Figure 1.**
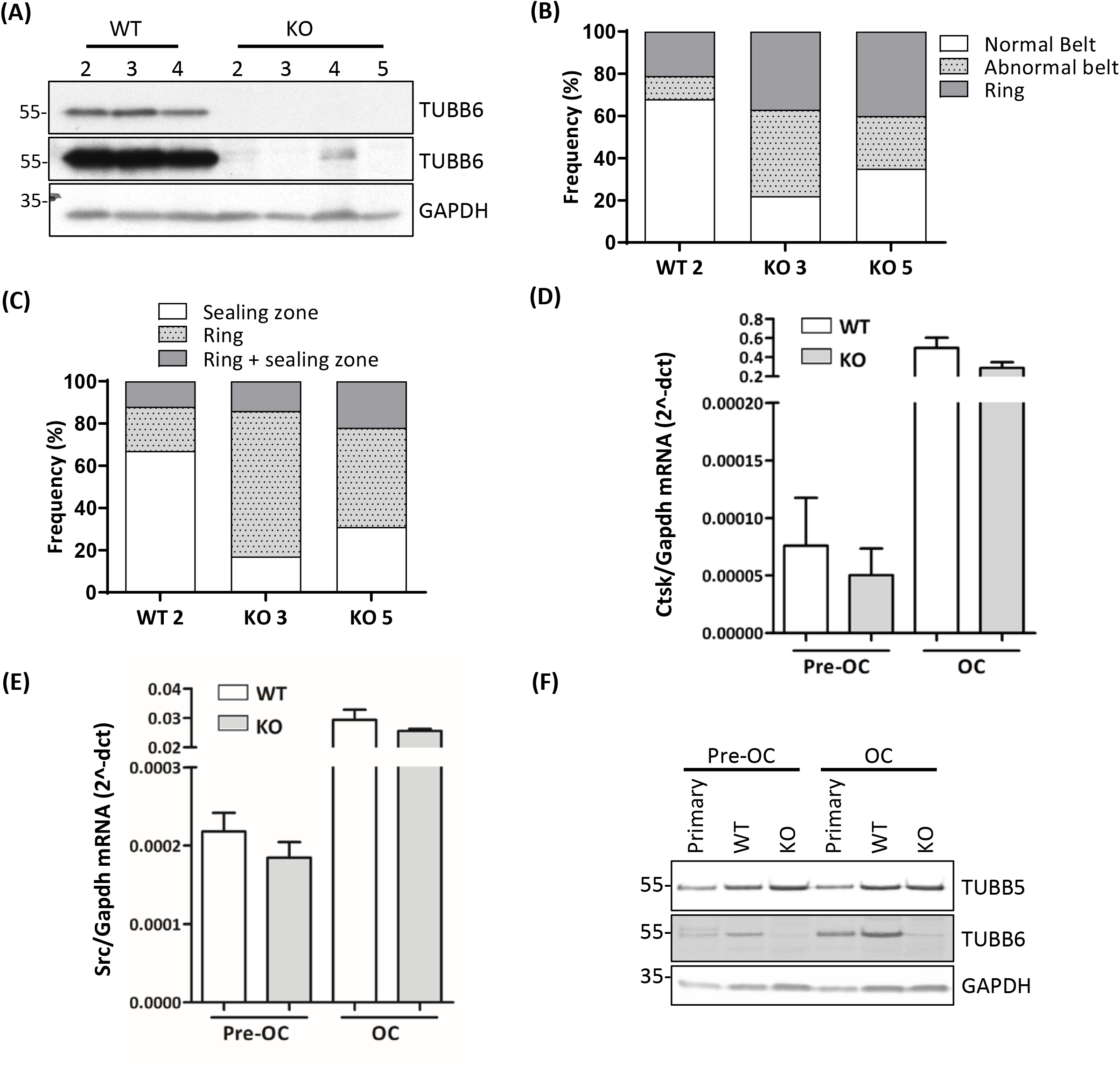
(A) Representative immunoblot showing TUBB6 and GAPDH expression in WT and Tubb6 KO RAW264.7 clones. Note in overexposed panel in the middle that clones 2 and 4 retain traces of TUBB6. (B) Bar graph showing the frequency of WT and KO osteoclasts seeded on glass and presenting podosome rings, normal or abnormal podosome belt, counting a total of 200 WT and KO osteoclasts in 2 independent experiments. (C) Bar graph showing the frequency of WT and KO osteoclasts seeded on ACC and presenting sealing zone and/or rings counting over 50 osteoclasts per clone. (D-E) Bar graphs showing the level of osteoclast characteristic Cathepsin K (D) and Src (E) mRNAs relative to Gapdh as determined by RT-PCR in WT or Tubb6 KO RAW264.7 cells (Pre-OC) or osteoclasts (OC). (F) Representative western blot showing TUBB5, TUBB6 and GAPDH expression in primary bone marrow macrophages (Pre-OC) and osteoclasts (OC) and in WT or Tubb6 KO RAW264.7 cells (pre-OC) or osteoclasts (OC).

**Supplementary Figure 2.**
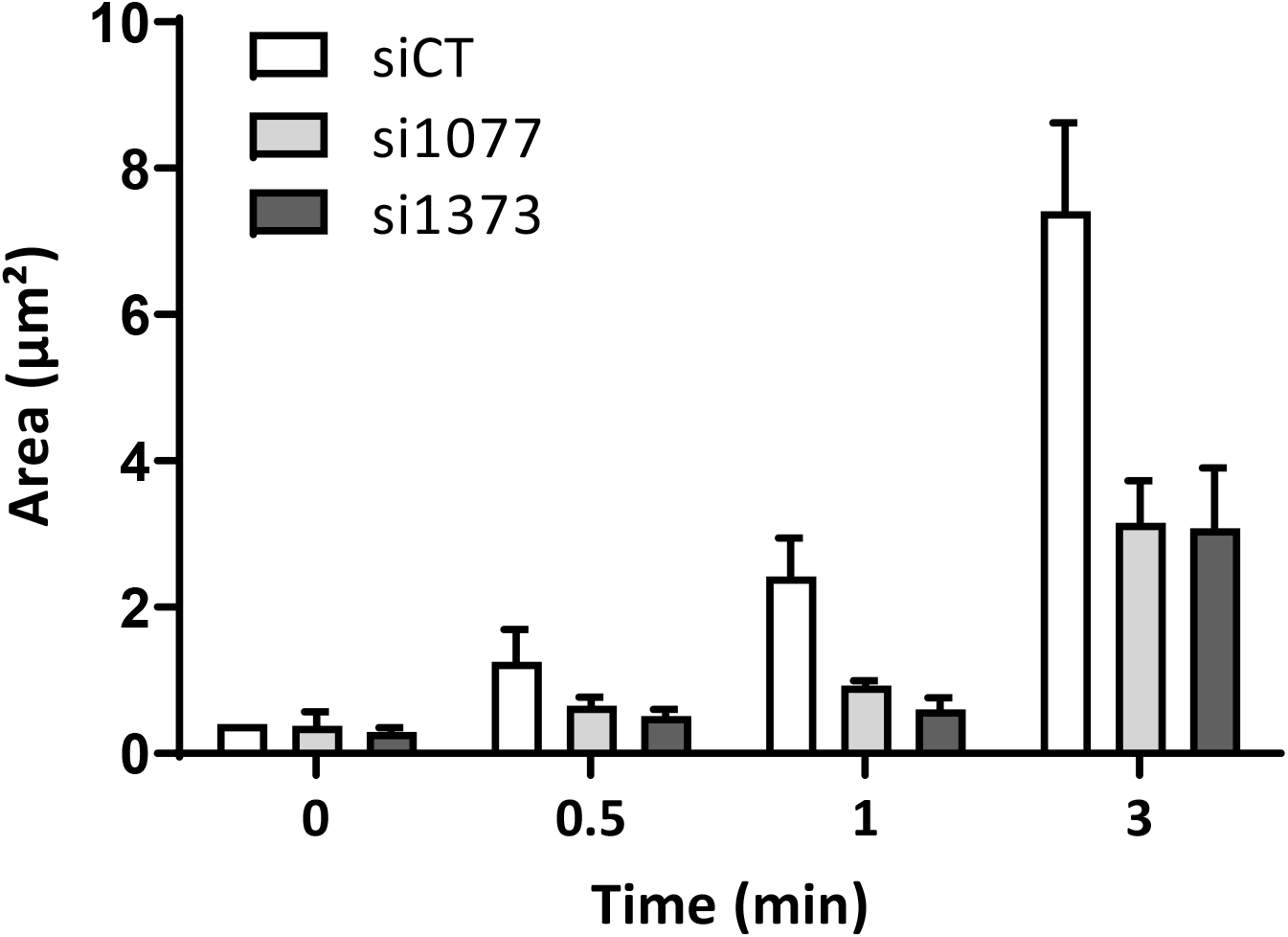
Bar graph showing mean and SEM α-tubulin aster area after nocodazole washout in osteoclasts transfected with luciferase control siRNA or Tubb6 siRNAs si1077 and si1373, measuring at least 20 asters per group were measured in 4 independent experiments.

**Supplementary Figure 3.**
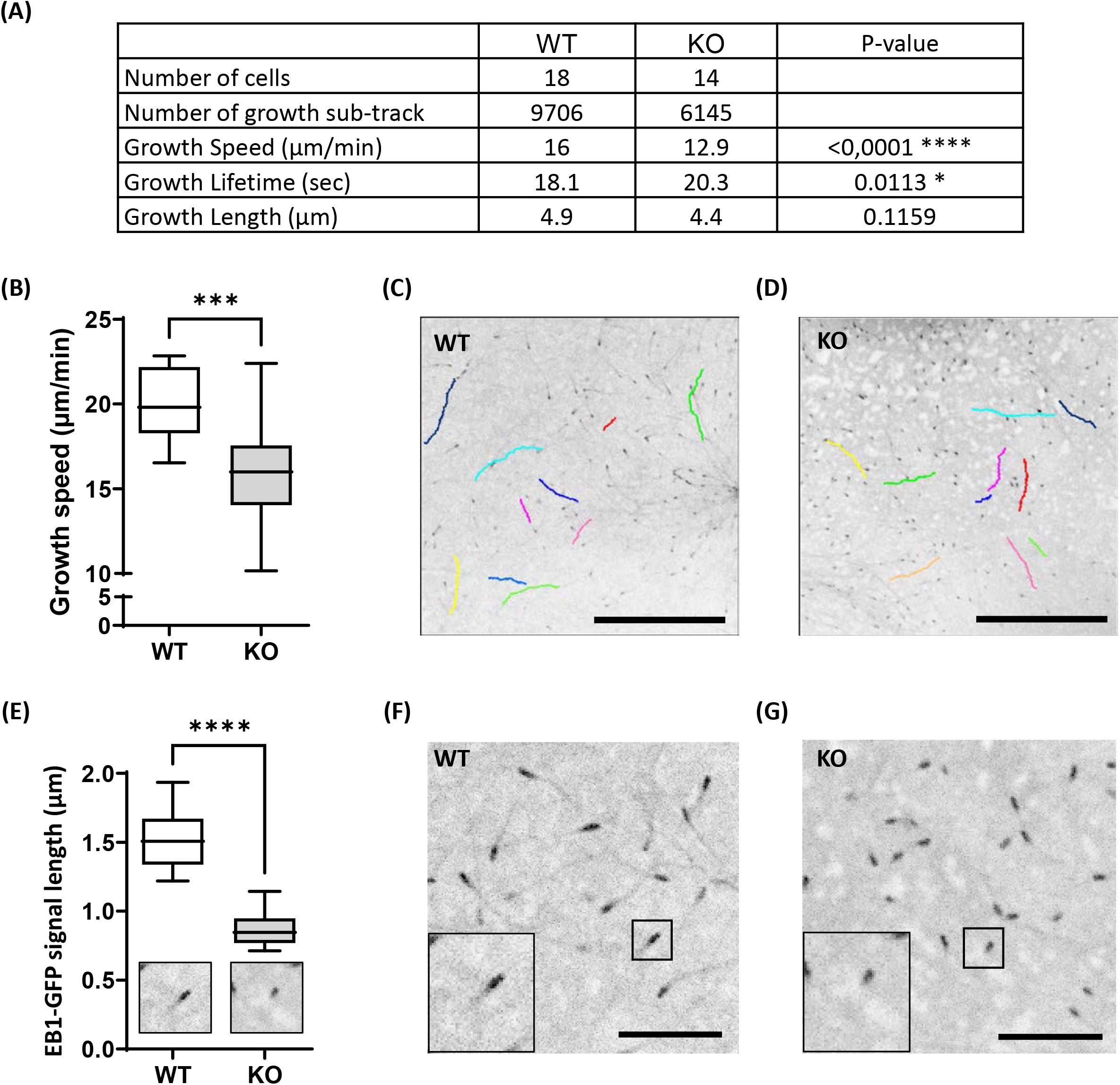
(A) Table showing the details of microtubule dynamics parameters from u-track analysis (B) Minimum to maximum boxplot showing microtubule growth speed determined in ImageJ 1.53c with MTrackJ in 15 WT and Tubb6 KO osteoclasts expressing EB1-GFP from three independent experiments and measuring 10-11 comets per osteoclasts; Mann Whitney test: ***p<0.0002. (C-D) Representative EB1-GFP comets tracks in WT (C) and Tubb6 KO (D) analyzed in B; scale bar: 10 μm. (E) Minimum to maximum boxplot showing the length of the EB1-GFP signal in 18 WT and Tubb6 KO osteoclasts per condition from three independent experiments and measuring 25 comets in each osteoclast to determine the average comet length per osteoclast; Mann Whitney test: ****p<0.0001. (F-G) Representative images of the EB1-GFP signal at the tip of microtubules in WT (F) and KO (G) osteoclasts analyzed in E; scale bar: 10 μm.

**Supplementary Figure 4.**
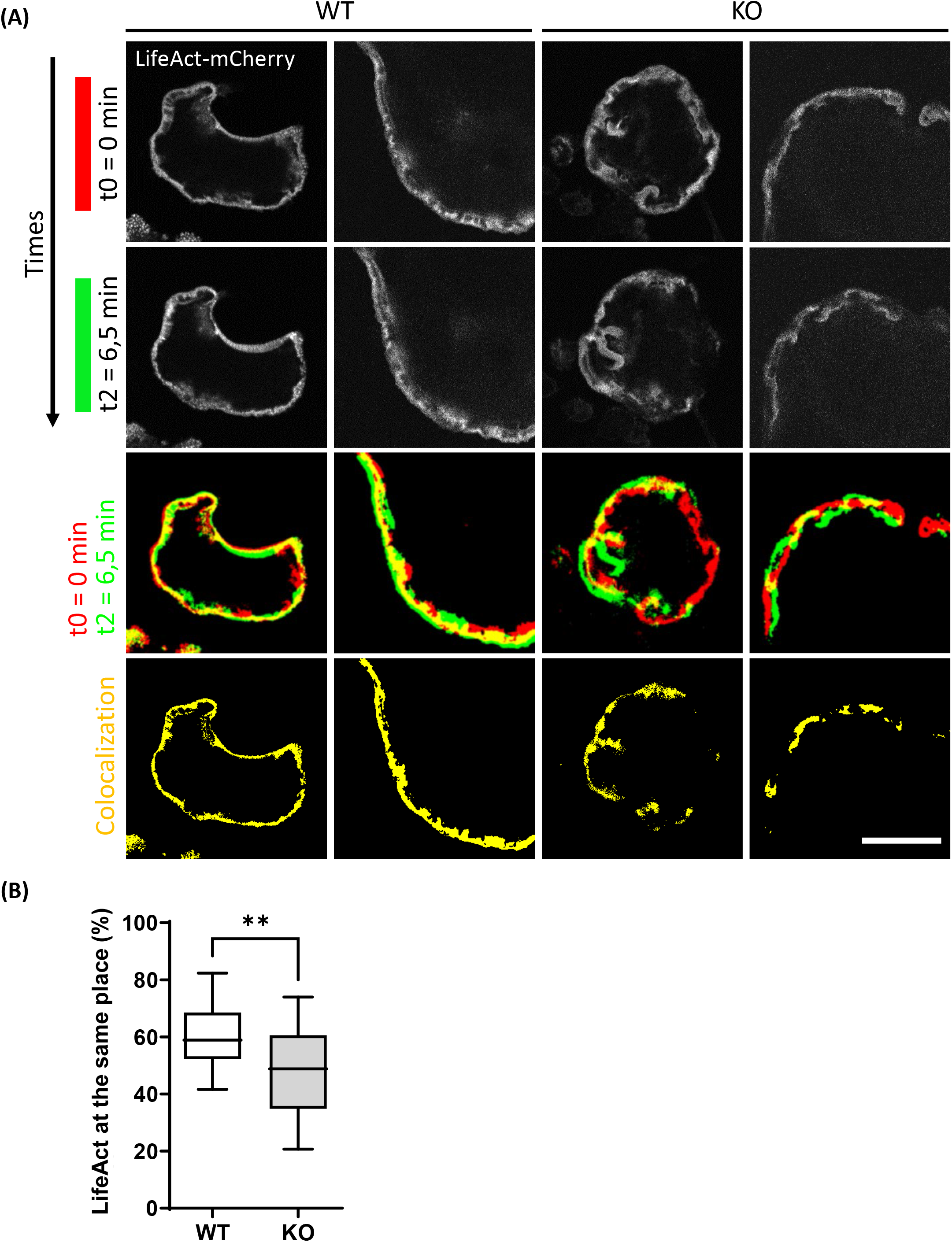
(A) Representative live confocal images of osteoclasts derived from WT or Tubb6 KO osteoclasts expressing LifeAct-mCherry and sitting on glass, showing the localization of LifeAct-mCherry signal at t0 = 0 minute (red) and t2 = 6.5 minutes (green), with the overlapping areas in yellow. Scale bar is 30 μm. (B) Minimum to maximum boxplot showing the percentage of LifeAct-mCherry signal at t0 that persists at the same position at t2 in WT or Tubb6 KO osteoclasts; in a total of 22 WT and 21 KO osteoclasts from 4 different experiments. Mann Whitney test: **p< 0.01.

*Supplementary Videos 1 and 2*

Detection and tracking of EB1-GFP expressing osteoclasts WT (video 1) or KO (video 2) using u-track software and settings described in Materials and Methods. Time is indicated as min:sec and frame is 68 μm.

*Supplementary Videos 3 and 4*

Live imaging of LifeAct-mCherry expressing WT (video 3) or KO (video 4) osteoclasts. Time is indicated as min:sec and frame is 90.5 μm.

*Supplementary Videos 5 and 6*

Live imaging of LifeAct-mCherry expressing WT (video 3) or KO (video 4) osteoclasts. Time is indicated as min:sec and frame is 16 μm.

